# Musical Insights into Protein Sequences and Functions via NMR Data

**DOI:** 10.1101/2025.03.09.642256

**Authors:** Zahra Moosavi-Movahedi, Parinaz Lordifard, Ramin Akhavijou, Kaveh Kavousi, Shohreh Ariaeenejad, Reza Behafarin, Amir M. Mortazavian, Abbas Shockravi, Ali Akbar Moosavi-Movahedi

## Abstract

This study explores an innovative technique for transforming rhythmic patterns found in nature into music, with a focus on protein sequences. For the past twenty years, various strategies have been developed for musical translation of biological sequences. This research presents a groundbreaking approach where music is derived from protein nano-scale structures using the atomic frequencies of amino acids extracted from NMR experiments. Each amino acid is assigned a unique musical note and duration, determined by its ^13^CNMR spectrum and relaxation period. Applying this method, music was composed from different protein sequences, including homologue β-globin, where a set theory-based comparative analysis was conducted. This novel approach opens up a new auditory dimension for understanding protein functions.

## 1. Introduction

Various studies have shown that rhythmic oscillations in natural phenomena can be transformed into tangible music for humans through diverse methods ^1–7^.

In ancient times, the relationship between sounds, mathematics, and physics was well-recognized. For example, the equations governing the motion of a vibrating string that produces a musical note were identified by physicists. However, identifying natural frequencies and composing musical notes require collaboration between scientists and musicians. Protein-derived music appears to reflect many of the protein features, such as its biological function, due to the intrinsic features of the amino acids^8^.

Since the 80s, scientists have found that there are significant similarities between DNA sequences, genes, and protein sequences with electronic music ^9,10^. Some music scholars, such as Alexjander, have conducted research on this subject ^11,12^. In Alexjander’s view, nature-based music is a type of nature-based music and the vibratory patterns in it. The movement in both macro and micro nano-dimensions (e.g., cosmic phenomena versus molecular and atomic surfaces) suggests multiple layers of meaning and surprising phenomena throughout the world. In the music composed by Alexjander, electromagnetic frequencies derived from the molecular world are used. The data used by artists like her come from a variety of sources, including atomic spectrographs such as DNA and water, most of which are in the IR spectrum. However, the question seems to be whether the basic constituents of living things and other elements of nature are harmoniously structured. In other scientific endeavors, mathematical equations have been used to translate sequences into musical notes. These equations are adjusted based on the physical features of the nucleotides in codons ^13,14^. Codons are regions consisting of three consecutive nucleotides in DNA that are often translated into amino acids in the protein polypeptide chain.

Perhaps the first tangible attempt in this regard can be made by Deamer, a biologist known for his efforts to understand the origin of life ^11,12^. The result of the joint study by Deamer and Alexjander was a piece of music called Sequencia, published on Earth Day (April 22, 1990) that used natural frequencies in DNA to produce it^12^. A strong, short bond, such as hydrogen bonding, absorbs the light spectrum with a higher wavelength (the number of waves alternating in one centimeter), which means a higher IR frequency in the IR spectrum and therefore lower frequencies in music. In Sequencia music, the sole criterion for musical composition was the same frequencies, and more complex DNA sequences such as genes and pairwise configuration had no role in the sequence. The instrument used to extract these frequencies included a spectrophotometer and a source of IR radiation in the range of 600-4000 cm^-1^. For example, the C-H bond absorbs IR light with a wavenumber of 2900 cm^-1^. The frequency corresponding to the 2900 cm^-1^ wavenumber would be 8.7×10^13 Hz, which is beyond the audible range of human hearing (20 to 20,000 Hz). With 36 consecutive divisions of 2, lower octaves could be obtained to eventually reach the human auditory range at 1266 Hz.

Converting light into sound was another debatable issue. Light and sound transmission media are completely different. But perhaps our problem here is the relation between the frequencies, not theirs. Therefore, the sounds produced in this way can be considered to be a good representative of the intended link. Due to the type of bonds in each open pair, these numbers are converted to sound by a synthesizer microtone-generating electronic keyboard. In microtone music, the interval between consecutive sounds is shorter than the standard note and requires more high-quality instruments. In the later work, instead of considering each open couple individually, each codon attributes a single musical note, a chord, or even a short piece of music (phrase), and thus the corresponding music is compiled.

With the exception of DNA sequences, the question arises whether it is possible to transform protein structures that are biologically meaningful into pieces of music that have positive aesthetic features. One of the answers to this question is the research done by Shi et.al 2007^15^. In this study, the researchers’ viewpoint was to convert protein sequences to music fragments using Morse code. The short music fragments, each of which formed on the basis of 20 amino acids forming the protein, were used to compose a complete music related to a protein. Like the basic Morses that were used to convey a variety of messages. They have also attempted to create different genres in protein music by creating rhythms based on data obtained from protein sequences. Perhaps the most important innovation was the use of short pieces of music of unequal sizes for each amino acid which resulted in the addition of rhythm to the resulting music. This approach is similar to using Morse code to senda message. As such, each amino acid may be translated into a single musical note, a mixture of notes, or even a short piece of music.

Amino acids are key constituents of proteins and are considered recurring units in all living organisms. There are approximately twenty amino acids that are components of proteins. A number of researchers have been analyzing and studying the features of amino acids since 1820^16^. Since World War II, analytical methods for studying the features of these biological components have advanced significantly. Microbiological assays using a lactic acid plant microbial, chromatographic decomposition, ion-exchange chromatography, and in particular gas chromatography in the determination of formal compounds D and L ^17^, are examples of these methods. Acid or alkaline hydrolysis^18^ and fermentation are also important methods for determining and producing amino acids.

The physical and chemical features of each amino acid are most affected by the oppositely charged amino and carboxyl groups as well as in the asymmetric amino acid α-carbon atom. All amino acids, except glycine, have such a carbon atom that they produce isomeric enantiomeric phenomenon and racemization. Crystallography of 23 amino acids using X-rays proved that their crystals differ greatly. The Raman spectrum of amino acids was used to provide the first direct optical evidence for bipolar structure. IR studies showed a sub-region in the NH frequencies of neutral amino acids. But in general IR analyses do not provide an accurate representation of amino acids ^19,20^.

NMR spectroscopy can analyze molecular structures based on their chemical shifts and coupling constants. Some methods use the effects of asymmetric solvents or diastereomeric reactants on the nuclei of different atoms. Two molecules can be paired by reflection symmetry if they are partially homogenous. The ^13^CNMR spectra of these groups are more affected by paramagnetic displacements than the ^1^HNMR spectra. The nature of the groups after enzymatic translocation can be determined by the ^13^CNMR spectra of labeled methyl groups^21–23^. Theoretically, the thermal equilibrium of the spin system is distributed by resonant frequency irradiation. During this process, not only do the equilibrium population ratios change but also macroscopic magnetization occurs, causing cross-magnetic fields (Mx and My). The spin system returns to thermal equilibrium after the oscillations cease. This process is accomplished through two mechanisms: the first through longitudinal or spin-lattice equilibrium and the other through spin-spin or transverse equilibrium time. Rotational, seismic, and electronics are very slow. In some cases, the full time to reach equilibrium may be a few seconds, a few minutes, or even a few hours. This study shows the features of amino acids using carbon relaxation instead of hydrogen. The relaxation time (T1) of carbon nuclei depends on the size and bonding state of the molecules, while the T1 of hydrogen nuclei is similar for different molecules. The carbon chemical shifts and T1 value are useful for amino acid analysis. ^1^HNMR spectroscopy is another method, but it can be complex and inefficient due to signal overlap or noise. Other methods include chromatography, black carbon polymer composites, rotational fluorescence, fluorescence, ultraviolet, and colorimetry. These methods are based on the biological and chemical relevance of aliphatic amino acids ^24–28^. Chromatography is an effective method for amino acid analysis, but new techniques based on black carbon polymer composites, rotational fluorescence, fluorescence, ultraviolet, and colorimetry have been developed. NMR spectroscopy is the preferred method for most researchers, as it can identify and characterize amino acids and their derivatives. NMR spectroscopy studies the physico-chemical and structural features of amino acids ^29–39^.

In the present study, this issue has been explored using a musical approach derived from protein sequences and various perspectives on composition. We examined the musical differences between properly functioning homologue globin proteins. The extraction of NMR-based features and utilization of these features to generate music derived from protein sequences for the purpose of a comparative study of protein function was the main issue of this research project.

## 2. Materials and Methods

### 2.1. Choosing the desirable protein for studying and synthesizing the music

The β-globin proteins were selected for music synthesis and music sequence analysis. The choice of β-globin is due to its role as a crucial oxygen-carrying protein in hemoglobin, essential for human life and many other organisms. This protein’s role across various organisms which exist in diverse biological conditions and habitats, demonstrates its adaptability in structure and function to different environmental and critical factors like oxygen partial pressure, pH, and temperature.

Therefore, comparing the β-globin music of creatures living in different environmental conditions is highly significant ^40,41^ The β-globin function is very important in meeting the needs of animal metabolism due to changes in their environmental conditions^42–44^. Bravenitz, Konigsberg, and Hill determined the first globin structure of the hemoglobin amino acids by the Edman method. The globin chains are covalently linked via proximal histidine to the heme iron atom. Near histidine has two very important roles: 1. stabilizing the heme and reducing the probability of the hominin being separated from the globin and 2. impact on the reactivity and activity of heme iron by altering the elasticity of its heme-binding which in turn is based on the allosteric properties of hemoglobin ^45^. The prosthetic hemoglobin group (Heme) is made of a ferrous tetrapyrrole ring structure. These four pyrroles are linked together by means of a Methane bond. Each ring consists of 4 methyl groups, 2 vinyl groups, and 2 propionic acid residues and therefore belongs to the ferriprotoporphyrin class IX (Heme b). Pauling and Corey first proposed the concept of a second protein structure ^46^. They tested different polypeptide models to find a way to reach the conformation they had found through α-creatinine X-ray diffraction^47,48^. Polypeptide chains are stabilized by folding into α-helical forms or β-folded plates. A-helix is one of the second most common structures in proteins. The globin subunits only have a spiral structure of the α helix. Normally, when hemoglobin has four subunits, it contains 75-80% of the α helix, and therefore the globin is a protein with an all α structure. It is believed that the high percentage of regular α-helical structure in hemoglobin is a major factor in the high stability of this protein^49^. These spirals are positioned around the heme to prevent the heme from being exposed to excess water and also to provide a suitable envelope for oxygen binding ^50^.

### 2.2. NMR spectroscopy experiments on amino acids

In the present project, T1 was measured for amino acids using D_2_O as a solvent in an almost saturated solution. The Bruker-AVANCE.300 NMR device was used with the pulse t1irpg program (specifically for measuring T1 using Inversion Recovery). To achieve higher resolution, the number of scans of each amino acid varied depending on the solubility of each amino acid, and this number was higher for amino acids that were less soluble in D_2_O. Tyrosine amino acid was not soluble not only in D_2_O but also in other polar solvents such as DMSO, methanol, acetone-d_6_, and CDCl_3_. Due to the low solubility of L-Tyrosine, information about this amino acid has been excluded from studies in this study. The Recovery Delay Time (RDT) program is used to measure T1. A 180° pulse is applied at the beginning of the experiment for each amino acid and at the end, a 90° pulse is applied to observe and detect the resting time^51^

The database under study consisted of 313 training data and 385 protein domain test data in different tests using the ETKNN and LibSvm classification engine evaluated with the help of the K-Fold Cross Validation classification method^25,52,53^. These tests were performed once with data extracted from the features based on NMR spectroscopy and later using chemical-physical features extracted from the profile site for each protein. The results of these tests have been studied from different perspectives. The main purpose was to compare the results of classifying by using the ETKNN engine on the characteristics obtained by NMR spectroscopy with the results of this classification on physical-chemical features^54–64^. These tests were performed under the same conditions^65^.

### 2.3. From Protein Sequence to Music

Protein-Melody is a MATLAB application designed to transform protein sequences into musical compositions. This program utilizes NMR data from the carbon atoms of amino acids (Abbas Shockravi et al., n.d.), specifically focusing on α carbon frequencies to represent each amino acid. To convert these frequencies into musical notes, the program divides the frequency intervals by two, enabling users to select start and end points within a range of 1 to 11 octaves, aligning with the human auditory spectrum. Additionally, the NMR relaxation times of the carbon atoms are employed to determine note durations, and the program incorporates pauses and bar lines into the musical composition. The resulting data sequence is exported in Music-XML format, ensuring compatibility with music analysis and display software such as Sibelius. For more detailed insights into the program’s functionality and its pseudo-code, please refer to the supplementary file provided.

### 2.4. Forte’s Set Theory analysis of synthetic music for proteins

The analysis of musical notes based on the set theory was first presented by Alan Forte, one of the leading music theorists ^66^. The theory of sets in music primarily examines the relationships between ordered or unordered song categories. In the sorted category, most of the focus is on the melody. In this first analysis, the prime forms for the sequences of the notes are obtained, and then the music is analyzed accordingly. Using this method, we can provide a musical insight into the differences and similarities in two or more sequences of music and consequently the protein sequence. We obtain the prime forms of the protein studied based on the secondary structure of the protein. From the second structure, the protein is divided into different groups (α-helix, β-content, random-coil). So, this can be a good point to analyze groups based on the set theory approach.

## 3. Results

### 3.1. The results of music analysis using Forte’s Set Theory

This section provides a comparative analysis of synthetic music for β-globulin of bar-headed goose, human, and sablefish sequences. In all the proteins studied, each part of the secondary structure elements is considered as a legato. The Legato is a piece of music that is played continuously and unceasingly. The set theory first deals with relationships between ordered or unordered song categories. In the sorted category, most of the focus is on the melody.

### 3.2. β-globin sequence analysis

We first identified the prime forms of music synthesized from the β-globin protein in humans, geese, and fish, based on their respective secondary structures. These structures were categorized into α-helix, β-sheet, and coil groups, forming the foundation for analysis using a set theory approach. For clarity, Table 1 presents each group by its prime form, aiding in the identification of similar sounds across groups. From this point forward, the goal is to explain Table 1, covering all its rows from top to bottom. (In this study, the terms ‘rows,’ ‘prime forms,’ and ‘sets’ all refer to Table 1.)

**Table 1.** the presentation of the fundamental structure of each subject based on the secondary structures of β-globin.

|  | Human (hmn) | Goose (gse) | Anoplopoma (ano) | statement |
| --- | --- | --- | --- | --- |
| 1 | (0,1,2,7) | (0,1,2,7) | (0,1,2,3,7) | Hmn and gse are the subsets of ano. Fig 2,4 |
| 2 | (0,1,2,3,4,7,8) | (0,1,2,3,4,6) | (0,1,2,3,4,5,8) | Hmn and ano have Rp relation. Fig 2,4 |
| 3 | (0,4) | (0,4) | (0,2) |  |
| 4 | (0,1,2,4,5) | (0,1,2,3,5,6) | (0,1,2,3,4,5,7,8) | Hmn is the subset of ano. Fig3, 5 |
| 5 | (0,1,2,3,4,5) | (0,1,2,3,5) | (0,1,2,3,5) | Gse and ano are the subsets hmn. Fig 2-5 ,6 |
| 6 | (0,1,3,7) | (0,1,3,4,7) | (0,1,4) | Hmn and ano are the subsets of gse. Fig 2,6,7 |
| 7 | (0,1,4,8) | (0,1,2,4,5) | (0,1,2) | Ano is the subset of gse. Fig 3,7,8 |
| 8 | (0,3) | (0,3) | (0,3) |  |
| 9 | (0,1,2,3,4,7,8) | (0,1,2,3,4,5,7,8) | (0,1,2,3,4,5,7,8) | Hmn is the subset of gse and ano. |
| 10 | (0,1,2) | (0,1) | 0 | Fig 2-8 |
| 11 | (0,1,2,3,4,7) | (0,1,2,3,4,5) | (0,1,2,3,4,5,7,8) | Both hmn and gse are the subset of ano. Hmn and gse have Rp relation. |
| 12 | (0,1,6) | 0 * | (0,1,6) | Three subjects are the same. (14<sup>th</sup> set of gse) |
| 13 | (0,1,3,4,5,6,8) | (0,1) | (0,1,2,3,4,5,8) | Hmn and gse are the same and they are in Rp relation with ano (0,1,3,4,5,8). |
| 14 | (0,1,2,4,7) | (0,1,6) | (0,1,2,4,7) | Hmn and ano are the same. Gse is the subset of two others. |
| 15 | (0,1,4,5,6,8) | (0,1,3,4,5,6,8) | (0,1,2,3,4,5,7,8) | Gse is the subset of ano. |
| 16 | (0,1) | (0,1,2,4) | (0,1) |  |
| 17 |  | (0,2,3,4,5,8) |  |  |
| 18 |  | (0,1) |  |  |

\* A zero in this table indicates that a category contains no distinct songs, only repeated scores.

This study aims to compare the rows of Table 1 (3-4) both in written analysis and through visual representation.

In comparative analysis, it’s important to recognize the similarities among many set classes, which is a key focus of this study. Table 1’s first row illustrates this point, showing that the β-globin melodies in both humans and birds share a common set (0,1,2,7), which is a subset of the fish category. Here, the concept of a superset becomes relevant. In this study, a sequence set (0,1,2,3,4,5,7,8) that contains all amino acid sounds is referred to as a superset. In ^13^CNMR spectroscopy of amino acids, many produce identical sounds after mapping, resulting in only 8 notes out of the 12 musical notes (according to semitone theory). Therefore, a set containing all amino acid sounds is defined as a superset. Fish exhibit 4 supersets, while birds have only 1, and humans do not have any supersets.

As previously mentioned, series categories that share the same prime form produce similar sounds. An examination of the second human set shows its repetition in the ninth set (Figure 2). Another example, shown in Figure 3, illustrates the identical nature of the fourth human set and the seventh bird set.

**Figure 1.**
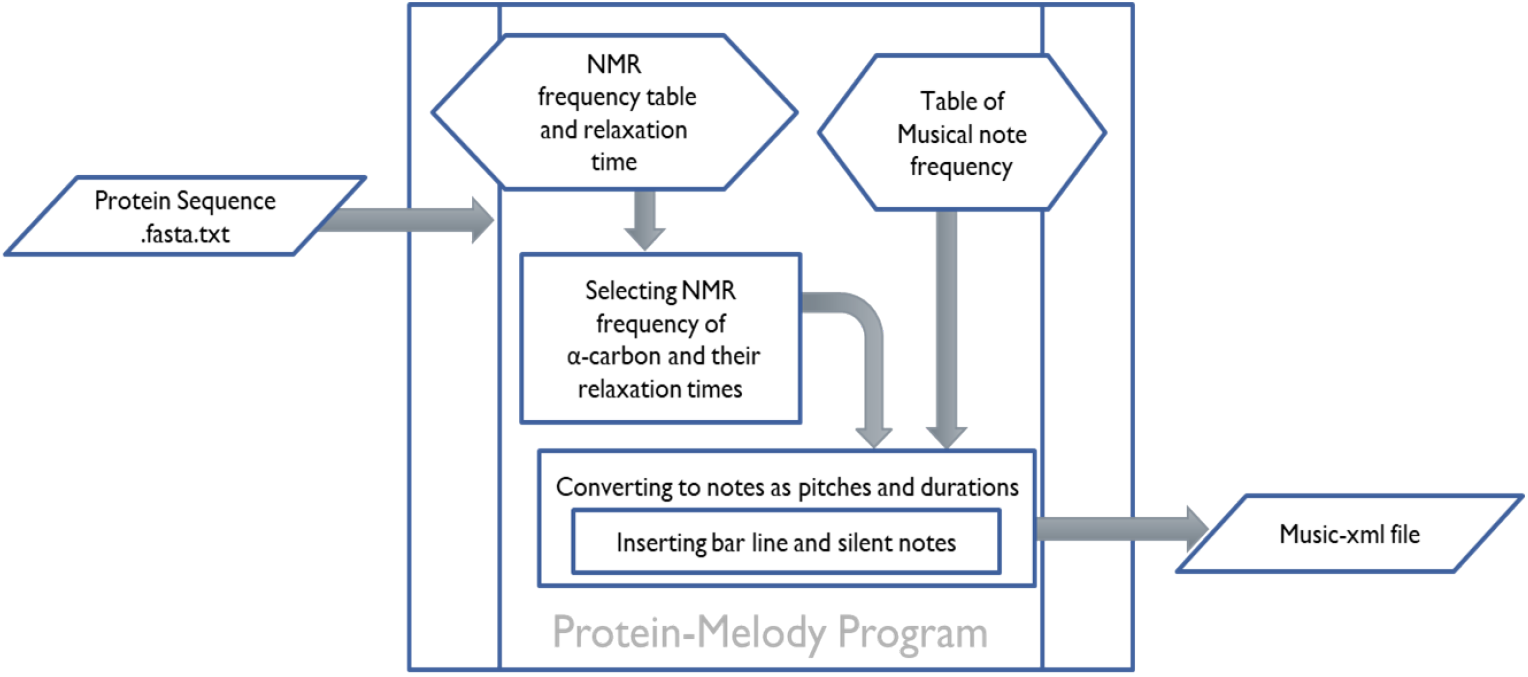
Workflow of the Protein-Melody program. This diagram illustrates the step-by-step process for transforming protein sequences into musical compositions, detailing the methods of data collection, note mapping based on NMR frequencies.

**Figure 2.**
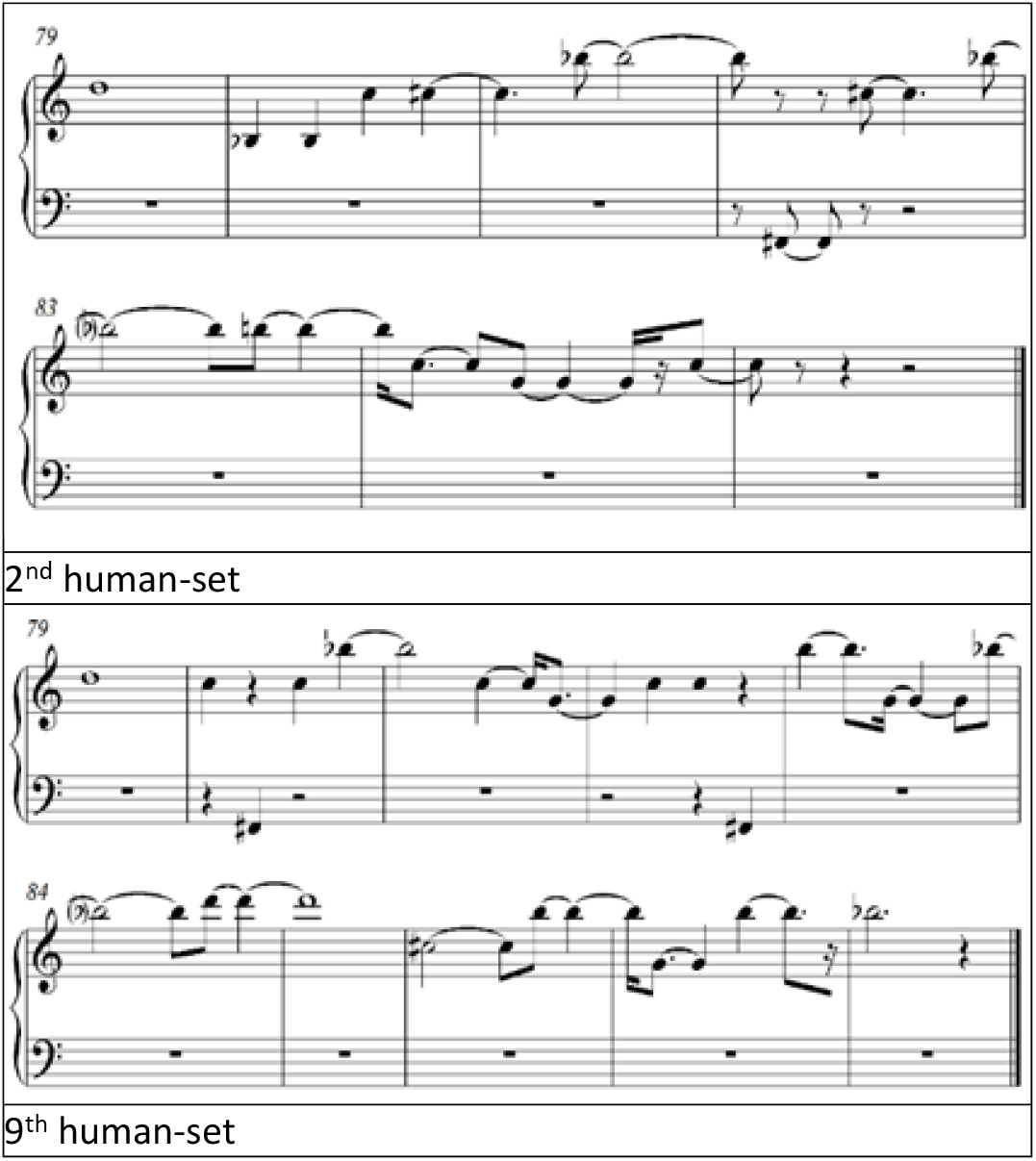
The second and ninth human classes share the same basic form. Despite differences in their melodies and structures, they are categorized as belonging to the same set.

**Figure 3.**
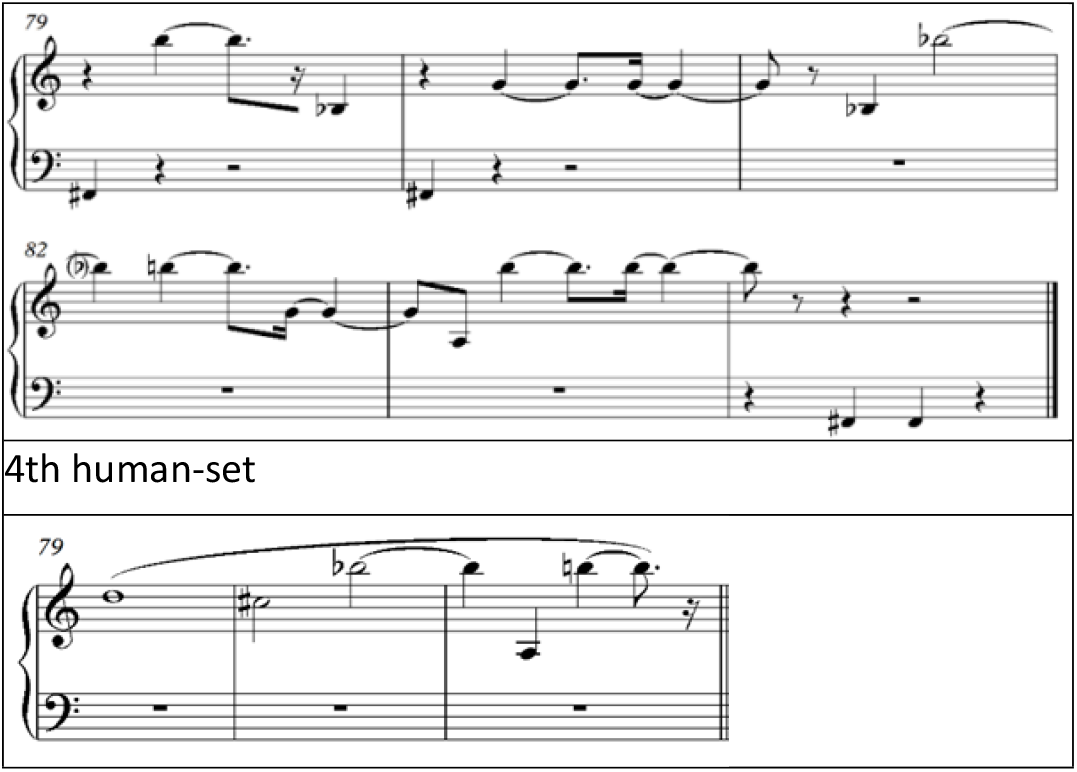
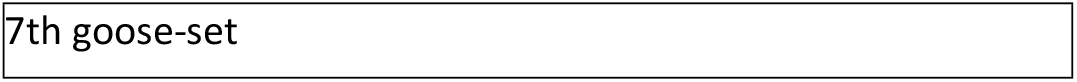
Comparison of two classes with identical sets from different β-globins (human and bird).

**Figure 4.**
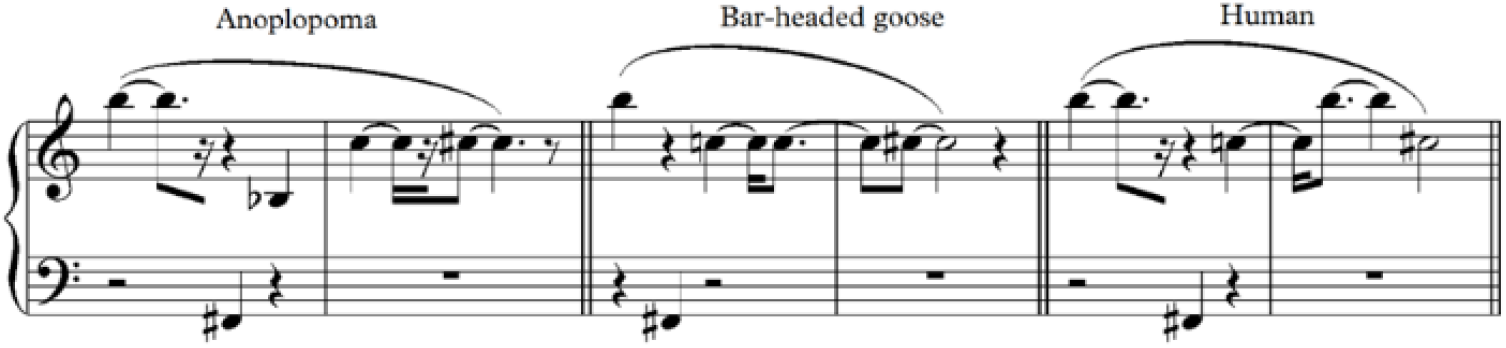
A visual comparison of musical sequences derived from the β-globin proteins of Anoplopoma (fish), bar-headed goose, and humans. The notation highlights the identical sets between humans and birds, both of which are subsets of the fish set. The difference is observed in the fourth note, where the bird’s double note becomes tonic in the human sequence.

**Figure 5.**
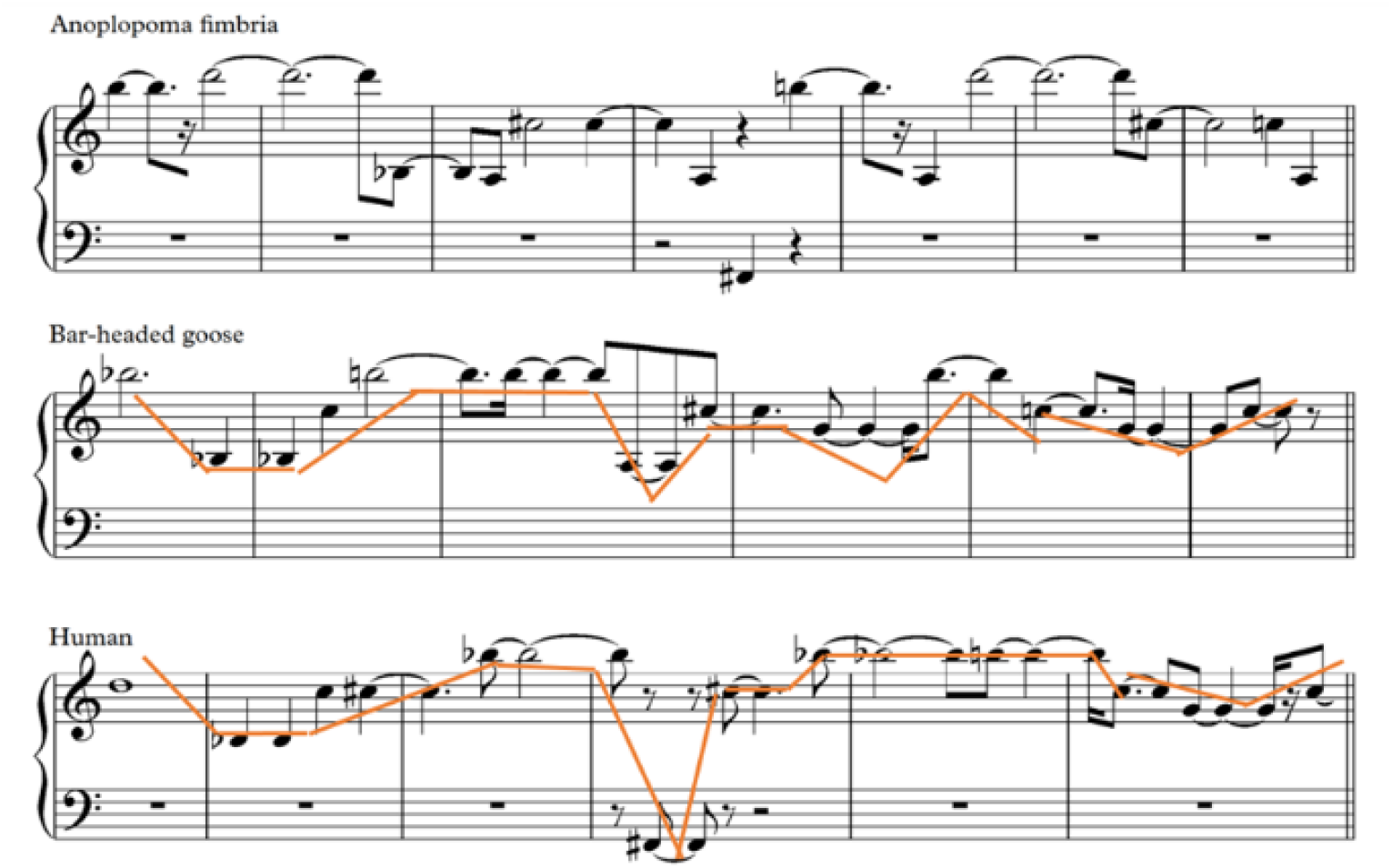
Comparison of melody designs among Anoplopoma fimbria (fish), bar-headed goose, and humans. The lines depict that humans and birds display more similar melodic designs compared to the fish. In the fifth row of Table 1, birds and fish share the prime form (0,1,2,3,5), with human subsets emphasizing the relationship between the human and bird sets.

In this figure, the human and bird samples show notable differences in musical elements such as melody, elasticity, and repetitive notes. However, they belong to the same category due to their identical prime forms: human (6,11,10,7,9) and bird (2,1,10,9,11). After normalizing, a set (1,2,3,5,6) is derived for humans and (9,10,11,1,2) for birds. Further analysis reveals that the bird’s 8-unit set (an 8-segment musical sequence) appears shifted, or alternatively, the human set is moved by 4 units. As a result, both sets align with the same prime form (0,1,2,4,5), creating a similar auditory experience for the listener.

The first row of Table 1 highlights the identical sets shared by humans and birds, both of which are subsets of the fish set. The key distinguishing factor is the inclusion of the element ‘3’ in the fish set. A discrepancy between humans and birds becomes apparent in the fourth note, where the double note in birds is replaced by a tonic in humans (moving from left to right in the figures). In the second row of the table, there is a clear similarity in the number of members across the three sets. Humans and fish both have sets containing seven members. Additionally, from the perspective of the Rp relationship —where Rp refers to the “relation-preserving” property in set theory, meaning that if there is a relation between elements of one set, and these elements are mapped to another set, the same relation is preserved (e.g., if set A is related to set B in a specific way, and each element of set A is mapped to an element in set B, the original relation between those elements remains valid)— they share the same subset (0,1,2,3,4,8). Furthermore, humans and birds exhibit similar melodic profiles.

The comparison of melody designs between humans, birds, and fish reveals that humans and birds display more similar designs, as illustrated by the lines in the figure, in contrast to the fish. In set theory analysis, single-member and two-member sets are typically not examined, except for cases like the progressive move represented in the tenth set. In the fifth row of Table 1, both birds and fish share the prime form (0,1,2,3,5), with human subsets reflecting this form. This emphasizes the similarity between the human and bird sets.

In the fifth row of Table 1, birds and fish share the same basic form, while human and bird melodies exhibit a notable similarity, with only one difference in their ninth note, as indicated by the arrow in the figure 6. Despite this, the fish melody still closely resembles the other two. Moving to row 6, it can be inferred that the human and fish sets are subsets of the bird set, which is derived from the union of these two sets. Consequently, the union of the human and fish sets (0,1,3,4,7) is contained within the bird category. As previously mentioned, the set theory approach exclusively focuses on musical intervals to identify the spatial similarity created by different compositions. To further understand the patterns in row 6, it is useful to examine the musical scores from this series, as shown in Figure 6, which highlights the significant resemblance between the melodies of humans and birds.

**Figure 6.**
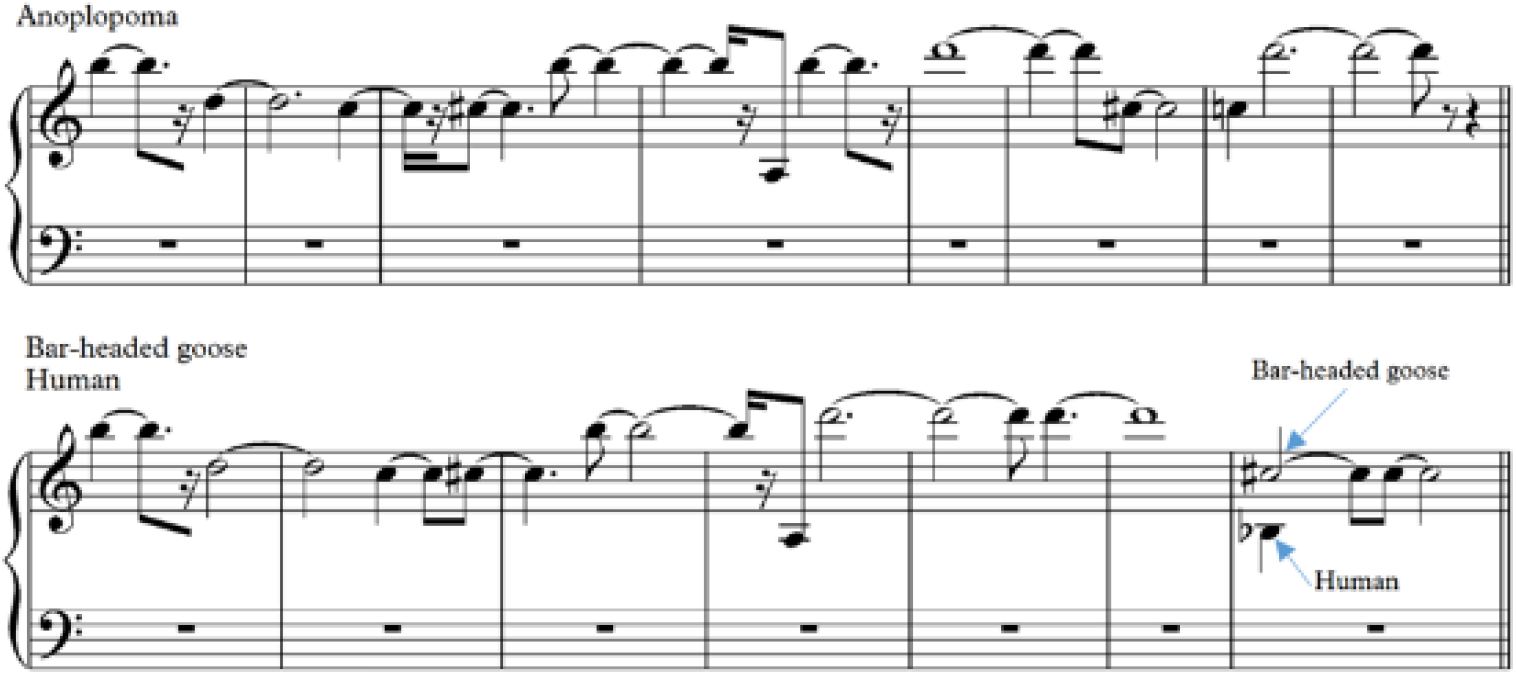
A comparative analysis of musical sequences from Anoplopoma, bar-headed goose, and humans in the fifth row of the Table 1. The figure illustrates the shared melodic structure between the bar-headed goose and humans, with specific transitions marked by arrows, highlighting their close relationship in contrast to the fish melody.

Figure 7 illustrates the seventh-row notes of the three proteins. Despite variations in their structural composition, all three proteins share a consistent melodic pattern, especially within the rectangular region. While F Sharp (indicated by the arrow) differs in the human series compared to the other two, the overall melodic design remains the same, with F Sharp functioning as a similar bass note. Due to the limited note options and their positions on amino acids, note A is the only alternative to represent the melody. The ninth complex, marking the beginning of the Heme region, is identical for both the fish and bird sets, with the human set being a subset due to its missing element. Unlike previous cases, the null set is not disregarded here due to a complementary process.

**Figure 7.**
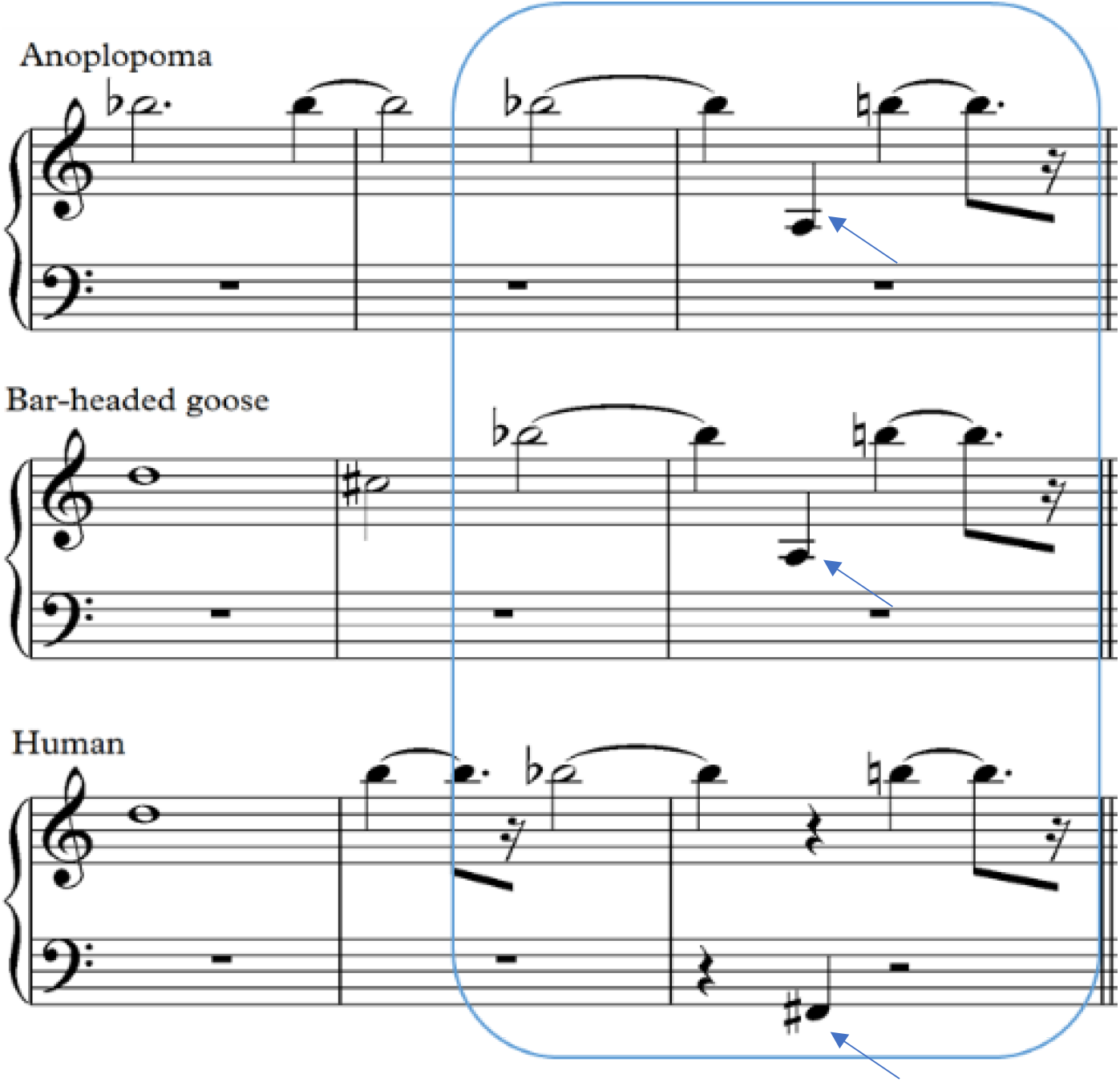
Comparison of seventh-row notes in Anoplopoma, Bar-headed Goose, and Human. The rectangular region highlights a shared melodic pattern. The arrows indicate variations in the F Sharp note. The ninth complex, associated with the Heme region, is identical in fish and birds but lacks an element in humans.

Figure 8 illustrates the tenth-row notes of the three subjects. The bird’s note is formed by adding two notes to the sides of the fish’s notes. However, the human’s first note deviates from the bird’s, suggesting a distinct pattern. This pattern change may indicate a divergence in the underlying biological processes or evolutionary pathways.

**Figure 8.**
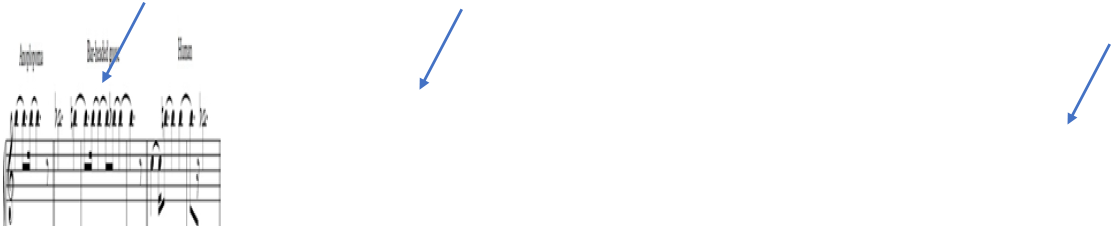
Comparison of tenth-row notes in Anoplopoma, Bar-headed Goose, and Human. The arrows highlight the differences in the notes, particularly the addition of notes in the bird and the deviation in the human first note.

In row 11, both the human and bird sets function as subsets of the fish set, sharing an Rp relationship with the same elements (0, 1, 2, 3, 4). Given the presence of two null sets in rows 12 and 13, it is logical to compare rows 12 through 16 of the human and fish series with rows 14 to 18. Notably, the prime form (0, 1, 6) is consistent across all three subjects.

Row 13 reveals that humans and birds are identical, sharing a common basic form (0, 1, 3, 4, 5, 8) in their Rp relationship with fish. Furthermore, row 14 highlights the similarity between the human and fish sets, both containing the set (0, 1, 2, 4, 7), while the bird set is a subset of the other two.

## 4. Discussion and Conclusion

The primary objective of this project was to develop a method for synthesizing music based on protein sequences. This approach facilitated the creation of unique musical compositions for each protein, representing either their complete or partial sequences, such as specific protein domains. The aim was to generate music that reflects the intrinsic properties of the proteins with minimal human intervention and variation in the musical composition. NMR spectroscopy served as the foundation for generating the notes, mapping the protein amino acids’ NMR frequencies to the human auditory range without altering their inherent musical features. As outlined in the Method section, the duration of each note corresponds to the relaxation time of the α carbon in the amino acids. Experts in the field of music have noted that the resulting compositions possess significant artistic meaning and interpretable elements.

This musical representation provides an aural basis for understanding differences between sequences that are closely related yet distinct. For instance, when creating music for homologous proteins with variations in their functional details, these differences become easily discernible and meaningful. In this study, we performed such a comparison on three β-globin sequences, which displayed notable differences in functionality influenced by environmental conditions like partial oxygen pressure. Set theory analysis effectively illustrated these distinctions, which were perceptible even to individuals without extensive musical knowledge. Moreover, applying Alan Forte’s systematic analysis method allows for a detailed exploration of the various regions within two or more proteins, revealing the musical elements of distinction and their influence on the overall composition. A significant outcome of this study is the potential to generalize scientific concepts and communicate them to society through artistic forms.

Additionally, the implications of this research extend to critical issues such as the structural and functional classification of proteins, predicting protein localization within cells, anticipating interactions with drugs, detecting protein complexes, and evaluating the medicinal properties of peptide sequences. The data related to resonant frequencies and the arousal times of atoms (the duration from excitation to return to the base state) were extracted from the ^13^CNMR spectrum of amino acids. These frequencies were then converted into sound frequencies corresponding to the 4th, 5th, and 6th octaves of the piano, with the note durations reflecting the excitation time of the atoms. This process resulted in melodies based on frequency and rhythm, which served as a foundation for further analysis.

Previous studies, such as those by Takahashi et al. (2007), employed a similar approach by assigning each of the 20 amino acids to specific notes within a reduced octave range and subsequently correlating them with chords to enhance the musical outcome. Similarly, Bywater and Middleton (2016) focused on selective proteins based on their NMR 3D structures, using the same frequency and rhythm principles to create music from different rhythms.

This study is groundbreaking in that it utilizes atomic excitation to produce musical melodies directly from the ^13^CNMR spectrum. The analysis of the generated melodies indicates observable variations in the sequence of notes associated with this music.

The main innovative goal of this research was to establish a scientific method rooted in the frequency features of amino acids at the atomic level, facilitating music production from the primary structural sequences of naturally occurring proteins on the nanoscale. By focusing on the NMR data from the α carbon atoms of amino acids, this research aims to address two fundamental questions: Does the music produced via this method reflect the evolutionary and natural selection processes from an aesthetic musical perspective? Or is the generated music merely a random sequence of notes? Future studies involving a broader range of proteins from different families could provide more comprehensive answers.

Another vital question concerns the potential to establish a meaningful connection between music derived from proteins and their functions. The present study demonstrates that changes in the amino acid sequence within protein-coding genes can lead to tangible alterations in the created music, making it meaningful for listeners. This suggests that variations in protein function may be detectable at the level of auditory representation, paving the way for new insights into protein functions through the lens of sound.

## 5. Acknowledgment

This work was supported by the Iran National Science Foundation (INSF), whose support is gratefully acknowledged.

## References

1. Wong, J.Y., McDonald, J., Taylor-Pinney, M., Spivak, D.I., Kaplan, D.L., and Buehler, M.J. (2012). Materials by design: Merging proteins and music. Nano Today 7, 488–495. 10.1016/j.nantod.2012.09.001.

2. Ghosh, R., Mishra, R.C., Choi, B., Kwon, Y.S., Bae, D.W., Park, S.C., Jeong, M.J., and Bae, H. (2016). Exposure to sound vibrations lead to transcriptomic, proteomic and hormonal changes in arabidopsis. Sci Rep 6. 10.1038/srep33370.

3. (a) Xing, Y., Xia, Y., Kendrick, K., Liu, X., Wang, M., Wu, D., Yang, H., Jing, W., Guo, D., and Yao, D. (2016). Mozart, Mozart Rhythm and Retrograde Mozart Effects: Evidences from Behaviours and Neurobiology Bases. Sci Rep 6. 10.1038/srep18744.

4. Bywater, R.P., and Middleton, J.N. (2016). Melody discrimination and protein fold classification. Heliyon 2. 10.1016/j.heliyon.2016.e00175.

5. Takahashi, R., and Miller, J.H. (2007). Conversion of amino-acid sequence in proteins to classical music: Search for auditory patterns. Genome Biol 8. 10.1186/gb-2007-8-5-405.

6. Schubert, M., Oschkinat, H., and Schmieder, P. (2001). MUSIC and aromatic residues: Amino acid type-selective 1H-15N correlations, III. Journal of Magnetic Resonance 153, 186–192. 10.1006/jmre.2001.2447.

7. Tay, N.W.N., Liu, F., Wang, C., Zhang, H., Zhang, P., and Chen, Y.Z. (2021). Protein music of enhanced musicality by music style guided exploration of diverse amino acid properties. Heliyon 7. 10.1016/J.HELIYON.2021.E07933/ATTACHMENT/83FA537D-BDD4-4489-9A3F-DD8F3987464E/MMC8.DOCX.

8. Dostrovsky, S. (1975). Early vibration theory: Physics and music in the seventeenth century. Arch Hist Exact Sci 14, 169–218. 10.1007/BF00327447.

9. Hayashi, K., and Munakata, N. (1984). Basically musical. Preprint, 10.1038/310096a0 https://doi.org/10.1038/310096a0.

10. Mallinckrodt, A.J., and Manning, P. (1988). Electronic and Computer Music. Leonardo 21. 10.2307/1578564.

11. Deamer, D. (2011). First life: Discovering the Connections between stars, cells, and how life began 10.5860/choice.49-1440.

12. Alexjander, S., and Deamer, D. (1999). The infrared frequencies of DNA bases: science and art. IEEE Eng Med Biol Mag 18, 74–79. 10.1109/51.752981.

13. Gena, P., Ph, D., and Strom, C. (1995). Musical Synthesis of DNA Sequences. XI Colloquio di Informatica Musicale.

14. Dunn, J., and Clark, M.A. (1999). Life Music: The Sonification of Proteins. Leonardo 32. 10.1162/002409499552966.

15. Shi, X.J., Cai, Y.Y., and Chan, C.W. (2007). Electronic music for bio-molecules using short music phrases. Leonardo 40. 10.1162/leon.2007.40.2.137.

16. Kinoshita, S. (1959). The Production of Amino Acids by Fermentation Processes. Adv Appl Microbiol 1. 10.1016/S0065-2164(08)70480-5.

17. Hasegawa, M., and Matsubara, I. (1975). Gas chromatographic determination of the optical purities of amino acids using N-trifluoroacetyl menthyl esters. Anal Biochem 63. 10.1016/0003-2697(75)90352-8.

18. Simpson, R.J., Neuberger, M.R., and Liu, T.Y. (1976). Complete amino acid analysis of proteins from a single hydrolysate. Journal of Biological Chemistry 251. 10.1016/s0021-9258(17)33637-2.

19. Herlinger, A.W., Wenhold, S.L., Longd, T.V., and Herlinger, A.W. (1970). Infrared Spectra of Amino Acids and Their Metal Complexes. II. Geometrical Isomerism in Bis (amino acidato)copper(II) Complexes. J Am Chem Soc 92. 10.1021/ja00725a015.

20. Pearson, J.F., and Slifkin, M.A. (1972). The infrared spectra of amino acids and dipeptides. Spectrochim Acta A 28. 10.1016/0584-8539(72)80220-4.

21. Purchase, R. (1987). NMR in chemistry: A multinuclear introduction. By W. Kemp. Macmillan Education Ltd, Hampshire, 1986. pp. xiii + 240. £25.00. ISBN 0-333-37291-3. Preprint, 10.1016/0278-6915(87)90135-9 https://doi.org/10.1016/0278-6915(87)90135-9.

22. Hill, R.K., Yan, S., and Arfin, S.M. (1973). Stereochemistry of valine and isoleucine biosynthesis. III. Enzymic discrimination between diastereotopic enol faces in the dehydrase step of vline biosynthesis. J Am Chem Soc 95. 10.1021/ja00804a048.

23. Baldwin, J.E., Löliger, J., Rastetter, W., Neuss, N., Huckstep, L.L., and La Higuera, N. De (1973). Use of Chiral Isopropyl Groups in Biosynthesis. Synthesis of (2RS5,3S)-[4-13 C]Valine. Preprint, 10.1021/ja00792a055 https://doi.org/10.1021/ja00792a055.

24. Buryak, A., and Severin, K. (2005). A chemosensor array for the colorimetric identification of natural amino acids. J Am Chem Soc 127, 3700–3701. 10.1021/JA042363V/SUPPLffFILE/JA042363VSI20050209_075710.PDF.

25. Lei, X., Liu, L., Chen, X., Yu, X., Ding, L., and Zhang, A. (2010). Pattern-based recognition for determination of enantiomeric excess, using chiral auxiliary induced chemical shift perturbation NMR. Org Lett 12, 2540–2543. 10.1021/OL100773S/SUPPL_FILE/OL100773S_SI_001.PDF.

26. Uccello-Barretta, G., Vanni, L., Berni, M.G., and Balzano, F. (2011). NMR enantiodiscrimination by pentafluorophenylcarbamoyl derivatives of quinine: C10 versus C9 derivatization. Chirality 23, 417–423. 10.1002/CHIR.20945.

27. Wright, A.T., Anslyn, E. V., and McDevitt, J.T. (2005). A differential array of metalated synthetic receptors for the analysis of tripeptide mixtures. J Am Chem Soc 127, 17405–17411. 10.1021/JA055696G.

28. Silverstein, R.W., and Bassler, G.C. (1962). Spectrometric identification of organic compounds. J Chem Educ 39, 546–553. 10.1021/ED039P546.

29. Frantz Folmer-Andersen, J., Kitamura, M., and Anslyn, E. V. (2006). Pattern-based discrimination of enantiomeric and structurally similar amino acids: An optical mimic of the mammalian taste response. J Am Chem Soc 128, 5652–5653. 10.1021/JA061313I/SUPPL_FILE/JA061313ISI20060327_092158.PDF.

30. Maier, N.M., Franco, P., and Lindner, W. (2001). Separation of enantiomers: needs, challenges, perspectives. J Chromatogr A 906, 3–33. 10.1016/S0021-9673(00)00532-X.

31. Severin, E.J., Sanner, R.D., Doleman, B.J., and Lewis, N.S. (1998). Differential detection of enantiomeric gaseous analytes using carbon black-chiral polymer composite, chemically sensitive resistors. Anal Chem 70, 1440–1442. 10.1021/AC970757H.

32. Nieto, S., Dragna, J.M., and Anslyn, E. V. (2010). A facile circular dichroism protocol for rapid determination of enantiomeric excess and concentration of chiral primary amines. Chemistry - A European Journal 16. 10.1002/chem.200902650.

33. Liu, H.L., Hou, X.L., and Pu, L. (2009). Enantioselective precipitation and solid-state fluorescence enhancement in the recognition of α-hydroxycarboxylic acids. Angewandte Chemie - International Edition 48. 10.1002/anie.200804538.

34. Shabbir, S.H., Joyce, L.A., Da Cruz, G.M., Lynch, V.M., Sorey, S., and Anslyn, E. V. (2009). Pattern-based recognition for the rapid determination of identity, concentration, and enantiomeric excess of subtly different threo diols. J Am Chem Soc 131. 10.1021/ja904545d.

35. Shabbir, S.H., Regan, C.J., and Anslyn, E. V. (2009). A general protocol for creating high-throughput screening assays for reaction yield and enantiomeric excess applied to hydrobenzoin. Proc Natl Acad Sci U S A 106. 10.1073/pnas.0809530106.

36. Bovey, F.A., and Tiers, G. V.D. (1959). Proton N.S.R. Spectroscopy. V. Studies of Amino Acids and Peptides in Trifluoroacetic Acid. J Am Chem Soc 81. 10.1021/ja01520a063.

37. Jardetzky, O., and Jardetzky, C.D. (1958). Proton magnetic resonance spectra of amino acids. J Biol Chem 233. 10.1016/s0021-9258(18)64769-6.

38. Sen, B., and Wu, W.C. (1969). Determination and identification of amino acids by thermometric titration and nmr spectroscopy. Anal Chim Acta 46. 10.1016/0003-2670(69)80039-5.

39. Nieto, S., Lynch, V.M., Anslyn, E. V., Kim, H., and Chin, J. (2008). High-throughput screening of identity, enantiomeric excess, and concentration using MLCT transitions in CD spectroscopy. J Am Chem Soc 130. 10.1021/ja803443j.

40. Qiu, Y., Maillett, D.H., Knapp, J., Olson, J.S., and Riggs, A.F. (2000). Lamprey Hemoglobin. Journal of Biological Chemistry 275. 10.1074/jbc.275.18.13517.

41. Perutz, M.F. (1978). Hemoglobin structure and respiratory transport. Sci Am 239. 10.1038/scientificamerican1278-92.

42. Giardina, B., Mosca, D., and De Rosa, M.C. (2004). The Bohr effect of haemoglobin in vertebrates: An example of molecular adaptation to different physiological requirements. Preprint, 10.1111/j.1365-201X.2004.01360.x https://doi.org/10.1111/j.1365-201X.2004.01360.x.

43. Berenbrink, M. (2006). Evolution of vertebrate haemoglobins: Histidine side chains, specific buffer value and Bohr effect. Respir Physiol Neurobiol 154. 10.1016/j.resp.2006.01.002.

44. Riggs, A. (1976). Factors in the evolution of hemoglobin function. Fed Proc 35, 2115–2118.

45. White, J.M. (1977). Haemoglobin structure and function: Its relevance to biochemistry and medicine. Preprint, 10.1016/0098-2997(77)90001-2 https://doi.org/10.1016/0098-2997(77)90001-2.

46. Pauling, L., Corey, R.B., and Branson, H.R. (1951). The structure of proteins; two hydrogen-bonded helical configurations of the polypeptide chain. Proc Natl Acad Sci U S A 37. 10.1073/pnas.37.4.205.

47. Dickerson, R.E., Kendrew, J.C., and Strandberg, B.E. (1961). The crystal structure of myoglobin: Phase determination to a resolution of 2 Å by the method of isomorphous replacement. Acta Crystallogr 14. 10.1107/s0365110×61003442.

48. Perutz, M.F., Muirhead, H., Cox, J.M., Goaman, L.C.G., Mathews, F.S., McGandy, E.L., and Webb, L.E. (1968). Three-dimensional fourier synthesis of horse oxyhaemoglobin at 2.8 Å Resolution : ((I) X-ray analysis. Nature 219. 10.1038/219029a0.

49. Nelson David L. (David Lee), 1942-. (2005). Lehninger principles of biochemistry. New York :W.H. Freeman,

50. Liong, E.C., Dou, Y., Scott, E.E., Olson, J.S., and Phillips, G.N. (2001). Waterproofing the heme pocket: Role of proximal amino acid side chains in preventing hemin loss from myoglobin. Journal of Biological Chemistry 276. 10.1074/jbc.M008593200.

51. Shockravi, A., Kavousi, K., Rezania, J., Jafari, R., Norouzi Beirami, M.H., Ariaeenejad, S., Moosavi-Movahedi, Z., Maghami, P., Mortazavian, A.M., and Moosavi-Movahedi, A.A. (2017). Time–frequency approach in the cluster assignment of amino acids based on their NMR profiles. Journal of the Iranian Chemical Society 14, 2221–2228. 10.1007/S13738-017-1158-1/FIGURES/2.

52. Ding, C.H.Q., and Dubchak, I. (2001). Multi-class protein fold recognition using support vector machines and neural networks. Bioinformatics 17. 10.1093/bioinformatics/17.4.349.

53. Yong-Sheng Ding, Tong-Liang Zhang, and Kuo-Chen Chou (2007). Prediction of Protein Structure Classes with Pseudo Amino Acid Composition and Fuzzy Support Vector Machine Network. Protein Pept Lett 14. 10.2174/092986607781483778.

54. Dubchak, I., Muchnik, I., Holbrook, S.R., and Kim, S.H. (1995). Prediction of protein folding class using global description of amino acid sequence. Proc Natl Acad Sci U S A 92. 10.1073/pnas.92.19.8700.

55. Hopp, T.P., and Woods, K.R. (1981). Prediction of protein antigenic determinants from amino acid sequences. Proc Natl Acad Sci U S A 78. 10.1073/pnas.78.6.3824.

56. Altschul, S.F., Madden, T.L., Schäffer, A.A., Zhang, J., Zhang, Z., Miller, W., and Lipman, D.J. (1997). Gapped BLAST and PSI-BLAST: A new generation of protein database search programs. Preprint, 10.1093/nar/25.17.3389

57. Schäffer, A.A., Aravind, L., Madden, T.L., Shavirin, S., Spouge, J.L., Wolf, Y.I., Koonin, E. V., and Altschul, S.F. (2001). Improving the accuracy of PSI-BLAST protein database searches with composition-based statistics and other refinements. Nucleic Acids Res 29. 10.1093/nar/29.14.2994.

58. Kaur, H. (2003). Prediction of beta-turns in proteins from multiple alignment using neural network. Protein Science 12. 10.1110/ps.0228903.

59. Kaur, H., and Raghava, G.P.S. (2003). A neural-network based method for prediction of γ-turns in proteins from multiple sequence alignment. Protein Science 12. 10.1110/ps.0241703.

60. Laurila, K., and Vihinen, M. (2011). PROlocalizer: Integrated web service for protein subcellular localization prediction. Amino Acids 40. 10.1007/s00726-010-0724-y.

61. Rashid, M., Saha, S., and Raghava, G.P.S. (2007). Support Vector Machine-based method for predicting subcellular localization of mycobacterial proteins using evolutionary information and motifs. BMC Bioinformatics 8. 10.1186/1471-2105-8-337.

62. Xie, D., Li, A., Wang, M., Fan, Z., and Feng, H. (2005). LOCSVMPSI: A web server for subcellular localization of eukaryotic proteins using SVM and profile of PSI-BLAST. Nucleic Acids Res 33. 10.1093/nar/gki359.

63. Cover, T.M., and Hart, P.E. (1967). Nearest Neighbor Pattern Classification. IEEE Trans Inf Theory 13. 10.1109/TIT.1967.1053964.

64. Zouhal, L.M., and Denoeux, T. (1998). An evidence-theoretic k-NN rule with parameter optimization. IEEE Transactions on Systems, Man and Cybernetics Part C: Applications and Reviews 28. 10.1109/5326.669565.

65. Kavousi, K., Moshiri, B., Sadeghi, M., Araabi, B.N., and Moosavi-Movahedi, A.A. (2011). A protein fold classifier formed by fusing different modes of pseudo amino acid composition via PSSM. Comput Biol Chem 35. 10.1016/j.compbiolchem.2010.12.001.

66. Benjamin, W.E., and Forte, A. (1974). The Structure of Atonal Music. Perspectives of New Music 13. 10.2307/832373.

